# Thermo-Flux: generation and analysis of comprehensive thermodynamic-stoichiometric metabolic network models

**DOI:** 10.1101/2025.11.20.689566

**Authors:** Edward N. Smith, Nathan Fargier, José Losa, Matthias Heinemann

## Abstract

Metabolic modelling, particularly through genome-scale stoichiometric models and flux balance analysis (FBA), has greatly advanced our understanding of metabolism. Yet, there is a continued quest to further improve the predictive capabilities of FBA. While thermodynamic constraints can allow for improved predictions, their addition to metabolic models has so far required cumbersome manual curation. To circumvent manual curation, we introduce ‘Thermo-Flux’, a semi-automated Python package, which converts stoichiometric metabolic network models into comprehensive thermodynamic-stoichiometric models for improved predictions. ‘Thermo-Flux’ enables (i) automated mass and charge balancing while considering physical and biochemical parameters, (ii) definition of transporter variants and Gibbs energies for membrane transport, (iii) robust handling of metabolites with unknown structures or Gibbs energies, and integrates (iv) recent advances in determining Gibbs energies and their respective uncertainties. To guide users in the conversion of stoichiometric models into comprehensive thermodynamic-stoichiometric models, we provide detailed instructions on how to use the ‘Thermo-Flux’ pipeline and include background information to enable appropriate modeling assumptions. By converting 87 stoichiometric models from the BiGG database and demonstrating improved flux predictions for a genome-scale yeast model (iMM904) converted into a thermodynamic-stoichiometric model, we showcase the applicability of ‘Thermo-Flux’. We expect ‘Thermo-Flux’ to support fundamental metabolic research and biotechnological applications. The package is available at https://github.com/molecular-systems-biology/thermo-flux along with tutorials.

## 2 Introduction

Metabolism is inherently complex and mathematical models provide a way to advance our understanding of its complexity. Genome-scale stoichiometric metabolic models combined with constraint-based approaches, most notably Flux Balance Analysis (FBA), have yielded valuable insights into metabolism [1, 2], ranging from the identification of essential metabolic functions and genes [3, 4] to the discovery of metabolic engineering targets [5, 6]. However, FBA can still also produce physiologically implausible flux predictions, highlighting the need for constraint-based approaches with improved flux prediction abilities [7].

The predictive power of FBA can be improved by complementing stoichiometric models with additional constraints, such as constraints on proteome allocation [8, 9, 10], or molecular crowding [11, 12]. Models can also be augmented with thermodynamics, which can constrain reaction directions and thus the feasible metabolic space [13, 14, 15], while also allowing the integration of measured metabolite concentrations [16, 17, 15]. The value of thermodynamic constraints in stoichiometric models was demonstrated by their ability to reveal fundamental cellular mechanisms, such as thermodynamic bottlenecks or metabolite channeling [18, 19, 20]. Recently, we went one step further and developed thermodynamic-stoichiometric models for *S. cerevisiae* and *E. coli* that incorporated comprehensive thermodynamics and accurate charge and proton balances for all reactions, including electron transport chains and transport reactions [21]. Used in combination with a constraint on the cellular Gibbs dissipation rate, these comprehensive models displayed improved predictive capabilities [21]. However, the manual effort required to convert stoichiometric models into such comprehensive thermodynamic-stoichiometric models is significant, which hampers their widespread use beyond the few commonly studied organisms.

To enable a broader application of thermodynamic constraints in metabolic modeling, automated tools are needed that reduce manual efforts during the development of such models. Tools such as multiTFA, pyTFA and matTFA, probabilistic thermodynamic analysis (PTA) and ThermOptCobra [22, 23, 19, 24] address some associated challenges, such as improved estimation of thermodynamic parameters and integration of metabolomics data. However, they lack several important features, such as automated charge and proton balancing and proper curation of transporter variants, accurate modeling of Gibbs reaction energies of transport processes, and guidance on how to handle the thermodynamics of atypical yet common reaction processes (e.g., biomass formation, electron transport chains or photochemical reactions).

To address these challenges, we introduce ‘Thermo-Flux’, a package designed to semi-automatically convert stoichiometric models into comprehensive thermodynamic-stoichiometric models. The key features of this package are: accurate charge and proton balancing based on physical and biochemical parameters of cellular compartments, automated definition of transporter variants and calculation of Gibbs reaction energies of transport processes, estimation of Gibbs energy of reactions with the component contribution method [25, 22], robust handling of metabolites with unknown structures or Gibbs formation energies, and the option to perform model regression to experimental data. In addition, we provide step-by-step instructions to guide users in the conversion of stoichiometric models into comprehensive thermodynamic-stoichiometric models. For each step, we describe the respective theoretical background and the assumptions made, illustrated by practical examples from a case study of the conversion of the yeast genome-scale metabolic network model (iMM904) [26]. Additional examples of the usage of ‘Thermo-Flux’ are available in the online documentation of the Python package (https://thermoflux-rtd.readthedocs.io/en/latest/). To demonstrate the applicability of ‘Thermo-Flux’, we converted 87 stoichiometric models in the BiGG database into thermodynamic-stoichiometric models with minimal manual curation. We conclude by showing the improved predictive capabilities of a fully parameterized thermodynamic-stoichiometric model. The package facilitates the development of comprehensive thermodynamic-stoichiometric models, which – through their improved predictive capability – will support fundamental and applied research questions on metabolism.

## 3 Overview workflow

The workflow of ‘Thermo-Flux’ to convert stoichiometric models into thermodynamic-stoichiometric models is divided into the following steps (Figure 1): (i) definition of physical and biochemical parameters (e.g. pH, ionic strength), (ii) definition of metabolites and their chemical species, (iii) calculation of metabolites’ Gibbs formation energies, (iv) identification of transported compounds, (v) application of pH-dependent charge and proton balancing, (vi) calculation of standard Gibbs energy of reactions including uncertainty, and (vii) formulation of the combined thermodynamic-stoichiometric solution space for constraints-based optimization. This last step also includes the possibility to fit model variables to experimental data such as extracellular reaction rates or metabolite concentrations. In the following, we provide instructions on how to perform each of these steps. In Appendix A, we also provide code snippets that illustrate relevant steps of the conversion of the iMM904 model.

**Figure 1.**
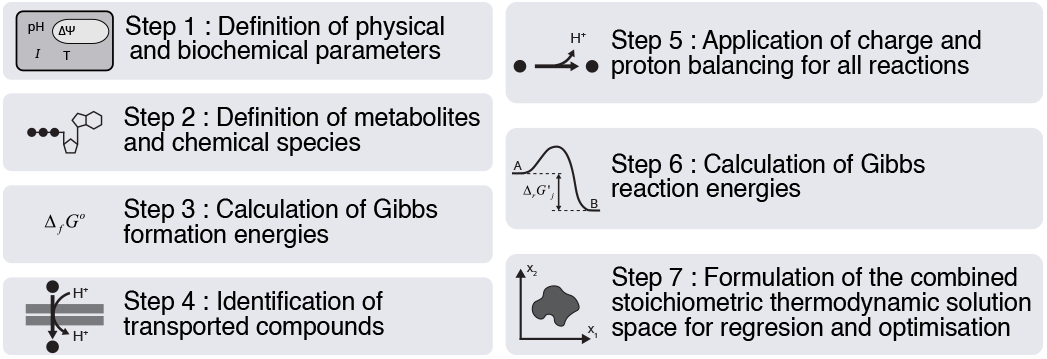
Overview of the different steps executed using ‘Thermo-Flux’ for converting a stoichiometric model into a comprehensive thermodynamic-stoichiometric model.

### Step 1. Definition of physical and biochemical parameters

The thermodynamics of biochemical reactions and transport processes is influenced by physical and biochemical parameters. For instance, pH, pMg and ionic strength determine the protonation and charge states of metabolites and their Gibbs formation energies. Membrane potential differences determine the Gibbs energy of transport reactions, and temperature influences all Gibbs reaction energies.

These parameters must be defined in the following manner: while temperature (*T*) takes a single value, others, such as pH, pMg, ionic strength (*I*) can take on different values in each of the subcellular compartments, as shown in Table 1. In addition, membrane potential differences (Δ*Φ*) are defined for pairs of compartments, i.e., the environments on either side of a membrane.

**Table 1.** Physical and biochemical parameters (and their units) used in the iMM904 case study. The code snippets used to set these parameters are in Appendix A, snippet 1.1.

| Parameter | Unit | Example values |
| --- | --- | --- |
| Potential hydrogen (pH) | Dimensionless | cytosol : 7, mitochondria : 7.4, extracellular space : 5 |
| Potential magnesium (pMg) | Dimensionless | all compartments : 14 |
| Temperature (T) | Kelvin (K) | T : 298 K |
| Ionic strength (I) | mol L<sup>-1</sup> (M) | all compartments : 0.25 M |
| Membrane potential difference ($\Delta\Phi$) | Volt (V) | plasma membrane : 0.06 V, mitochondrial membrane : 0.16 V |

Unless specified by the user, ‘Thermo-Flux’ uses default parameters for all subcellular compartments, i.e. pH = 7.0, pMg = 3.0, ionic strength = 0.25 M, membrane potential difference = 0 mV between two compartments. Yet, we advise users to always give preference to values reported for the specific organism and conditions of interest, as these values are important for realistic thermodynamic calculations. As reference, we provide physiological example values for the physical and biochemical parameters obtained from *E. coli, S. cerevisiae* and *Arabidopsis* (Supplementary table S1).

### Step 2. Definition of metabolites and chemical species

To be able to add proton and charge balances to the model and to calculate the Gibbs energies for reactions, we must first identify (i) the chemical structures of the metabolites in each reaction, and (ii) the chemical species that compose each metabolite and the proportions between the different species. To identify the structure of a metabolite, we use *eQuilibrator*, which stores structures for many biochemical compounds. To access these structures in *eQuilibrator*, we need to connect a metabolite, as provided in a reaction of a stoichiometric model, to the respective compound in *eQuilibrator*. This connection is achieved using various annotations that are typically provided in stoichiometric models. The annotations that can be used in ‘Thermo-Flux’ range from structural notations (InChI or SMILES) to databases identifiers, such as KEGG or BiGG (Figure 2a). Once each metabolite is linked to its corresponding compound in *eQuilibrator*, it is possible to query its chemical structure, as well as its acid and magnesium dissociation constants, i.e. *K*_*a*_ and *K*_*Mg*_.

**Figure 2.**
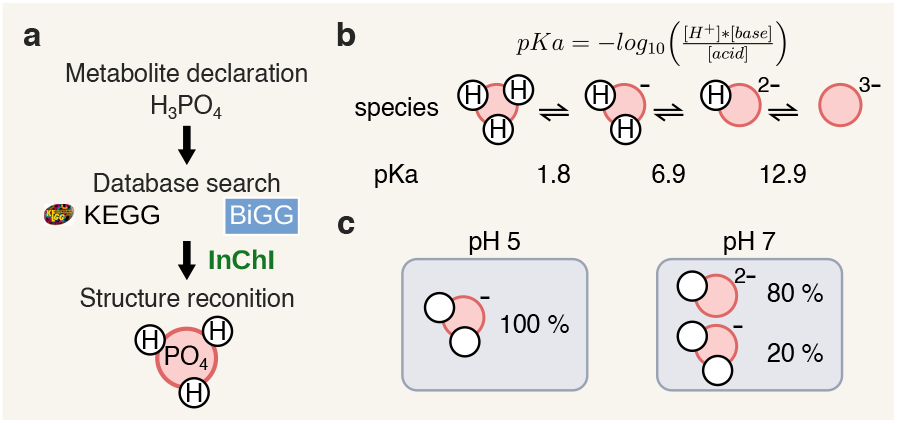
From metabolite definition to chemical species specification. a) From the definition of a metabolite using one of the multiple database annotations, the metabolite structure is obtained from the respective database or explicitly defined with its InChi notation b). From *pK*_*a*_ calculation based on ChemAxon and the pH, the eQuilibrator package is then used to calculate the chemical species distribution. The *pK*_*a*_ values of phosphate species in solution are shown. c) Given the proportions of each species of phosphate in pH 5 and in pH 7, the protonation and charge states vary in these two different conditions (2 protons and -1 charge at pH 5, 1.2 protons and –1.8 charge at pH 7)

In aqueous solutions, metabolites exist as a combination of multiple species differing in protonation and charge states. The proportion between these different species is defined by their acid dissociation constants and the physical and biochemical parameters (i.e., ionic strength and pH) (Figure 2b). The combination of these species is collectively referred to as a reactant, for which we determine the average protonation and charge states using its species distribution. When the proportions of each species are known, the average protonation and charge states of the reactant can be calculated. For example, inorganic phosphate (P_i_) can exist as species PO_4_^3-^, HPO_4_^2-^, H_2_PO_4_^-^ and H_3_PO_4_ (Figure 2b). At pH 7.0, P_i_ is composed of about 80% 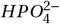 and 20% 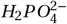 resulting in an average of 1.2 protons and an average charge of -1.8 for the reactant *P*_*i*_ (Figure 2c). Thermo-Flux automatically calculates the species distributions whenever a new metabolite is defined. The user can query the protonation and charge states of reactants and inspect the resulting species distribution estimated by ‘Thermo-Flux’ using the code snippets 2.1 and 2.2, respectively (Appendix A).

For metabolites whose structures cannot be encoded by InChI, such as complex organometallics (e.g. heme O) or metal ions, or for metabolites with unknown or ambiguous chemical structures, such as acyl carrier proteins, the protonation and charge states cannot be calculated. In these cases, we rely on literature references to select one species with a specific protonation and charge state that represents the average protons and charge of these metabolites. For example, for iMM904 we manually defined the charge of the redox couple ferricytochrome/ferrocytochrome, using the code snippet 2.3 (Appendix A).

Similarly to the equilibrium between species with different protonation states, an equilibrium can exist between species that form different complexes with Mg^2+^, with their abundances depending on their affinity to magnesium. With the same approach as described above, *K*_*Mg*_ values are used to estimate an average number of magnesium ions for a metabolite.

### Step 3. Calculation of Gibbs formation energies

The next step is to determine the standard Gibbs formation energies (Δ_*f*_ *G*^*°*^) for all metabolites. The Gibbs formation energy describes the Gibbs energy change required to form one mole of the metabolite from its constituent elements in their standard state (i.e., pH 7, concentration of 1 M). A way of estimating Δ_*f*_ *G*^*°*^ is to use the component contribution method (CCM). This method starts by decomposing compounds into constituting ‘components’ (i.e., smaller metabolites or chemical groups) with known Gibbs formation energies. The standard Gibbs formation energy of the metabolite is obtained by summing the standard Gibbs formation energies of its constituting ‘components’.

Using the CCM, we can obtain standard formation energies of reactants from *eQuilibrator*. However, as mentioned above, metabolites are present as combinations of species, which differ by their protonation states (step 2) and Gibbs formation energy. Specifically, the Gibbs formation energy of each successive protonation state of a metabolite can be expressed relative to the previous, less protonated form as

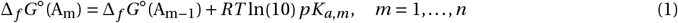

where *pK*_*a,m*_ is the acid dissociation constant for the protonation step from *A*_*m−*1_ to *A*_*m*_, where *A*_0_ would denote the fully deprotonated species.

Importantly, because the pH or ionic strength of a cell usually do not reflect the standard conditions, realistic thermodynamic calculations require that all Gibbs formation energies are corrected for the actual state of the cell. Specifically, a certain pH defined for a cell or compartment implies that the concentration of protons is set and remains constant. The Gibbs formation energies then need to be adjusted to this specified pH. In ‘Thermo-Flux’, these adjustments to cellular conditions are achieved by combining a Legendre transformation (correcting pH) with the extended Debye-Hückel equation (correcting ionic strength) [27], according to

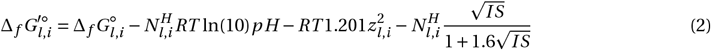

where *N*^*H*^ is the number of hydrogen atoms in the species, *I* is the ionic strength and *z* the charge of the species *l*. Then, the Gibbs formation energy of a metabolite is the “Boltzmann-weighted sum” of the transformed Gibbs formation energies of the constituent species [28] :

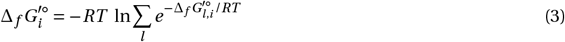

Here, 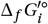 is the transformed Gibbs formation energy of metabolite *i, R* is the universal gas constant, and *T* the temperature. The Boltzmann weight for a species *l* of metabolite *i*, exp 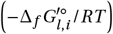, is determined by its transformed Gibbs energy of formation, which depends on the specified pH (Eq. 2) and its *pK*_*a*_ (Eq. 1). The fraction of this species *α*_*l*,*i*_ under the specified conditions is obtained by normalizing its Boltzmann weight by the sum of the Boltzmann weights of all species 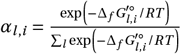.

Δ_*f*_ *G*^*′°*^ values are in principle condition-independent parameters. Yet, the component contribution method is based on experimental data and therefore not only returns estimates of these values, but also their respective uncertainties. These uncertainties can be cause by various sources, ranging from measurement errors in the determination of equilibrium constants to systematic errors embedded in assumptions made in the estimation process. Considering these uncertainties in thermodynamic-stoichiometric models avoids biasing the Gibbs energies towards a single set of values that could bear errors from these multiple sources.

After having defined the physical and biochemical parameters (step 1) and the metabolites of the model (step 2), we used the code snippet 3.1 (Appendix A) to calculate Δ_*f*_ *G*^*°*^, the transformed Δ_*f*_ *G*^*′°*^ and the corresponding uncertainties in Δ_*f*_ *G*^*°*^ for all metabolites of the iMM904 model.

#### Box 1

**Additional considerations for Gibbs formation energy calculations**

**Metabolites with unknown or non-decomposable structure**

Gibbs formation energies cannot be estimated for metabolites with an unknown structure or with a structure that cannot be decomposed by the component contribution method. For such metabolites, we assign a value of 0 kJ/mol, in combination with a large uncertainty of *±*3000 kJ/mol. This wide uncertainty range reflects the fact that the cellular Δ_*f*_ *G*^*′°*^ could lie anywhere within this range.

**Redox carriers**

For protein-based or membrane-bound redox carriers, such as cytochrome C, the Δ_*f*_ *G*^*′°*^ can be calculated from experimentally determined midpoint potentials using the Nernst equation for a redox half reaction,

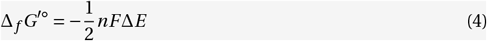

where *n* is the number of electrons transferred, *F* is the Faraday constant and Δ*E* is the redox reaction’s midpoint potential difference (in volt V).

**Photons**

The Gibbs formation energy of a photon is related to its wavelength *λ* by the equation

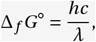

with *h* being Planck’s constant and *c* the speed of light in the growth medium.

**Biomass**

In stoichiometric models, biomass is also considered as a metabolite. To comprehensively describe the thermodynamics of the whole metabolic network including biomass formation, a Gibbs formation energy must also be specified for biomass. Values for Gibbs formation energies of biomass have been estimated based on the elemental composition of biomass and empirical formulas [29, 30]. These values must also be transformed based on the number of hydrogen atoms in the biomass empirical chemical formula. Transformation of the biomass metabolite’s Gibbs formation energy is important when applying a constraint on the rate of the cellular Gibbs energy dissipation as this value contributes to the total Gibbs energy dissipation which could be compared between cells with different biomass composition [21].

For special metabolites as illustrated in this box, users can manually define Gibbs formation energies and associated uncertainties in ‘Thermo-Flux’.

### Step 4. Identification of transported compounds

Transport reactions are processes that move metabolites across membranes. However, not all chemical species of a metabolite are necessarily transported, as transporter proteins can be specific to a particular species [31]. Yet, the protonation state of the transported species and its charge will determine the Gibbs energy change of the transport process (step 6) [32]. Therefore, to correctly describe the thermodynamics of a transport process, it is important to know the specifically transported species, as this species determines the net proton and charge movement across the membrane. The identification of the transported species can be based on biochemical data obtained from transport assays or inferred from transporter structures [33], if available.

Because this precise information is often not available, ‘Thermo-Flux’ uses the assumption that the chemical species with the highest abundance in the inner compartment is the one that is being transported. The process through which ‘Thermo-Flux’ determines the transported species, its protonation state and the net charge movement is the following. First, for any given transport process, metabolites that occur on both sides of the reaction equation are identified as being affected by the transport of its species across the membrane. Second, following our assumption, the species that has the highest abundance fraction in the inner compartment is selected as the one which is transported. Third, we calculate the protonation state of this species and the amount of charge that cross the membrane (Figure 3a). For example, considering the transport of phosphate from the extracellular space (pH 5) to the cytosol (pH 7) and applying the steps mentioned above, we (i) identify that phosphate is the transported metabolite, (ii) determine that the transported species is 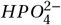, because it is the most abundant species in the inner compartment, and (iii) calculate that this transport process moves a species that has one proton and two negative charges.

**Figure 3.**
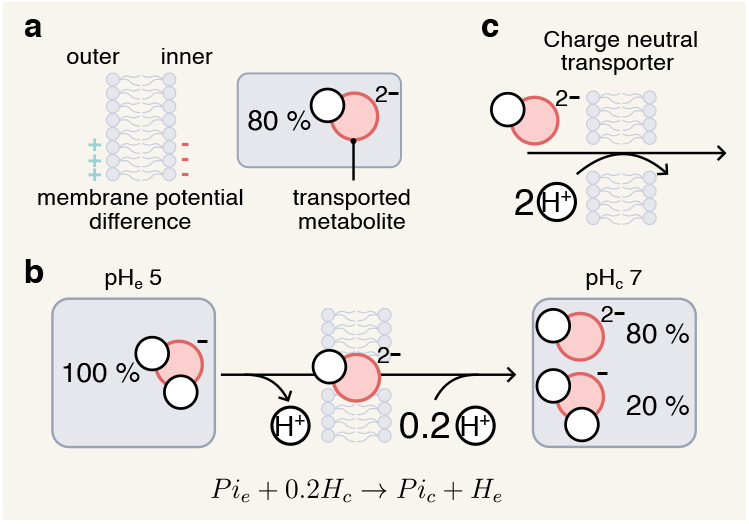
Identification of the transported compound a) The compartment with the lowest membrane potential is assumed to be the inner compartment. In this inner compartment, ‘Thermo-Flux’ identifies the major species of phosphate at pH 7; this species 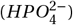 is assumed to be transported. b) In the uniport process of phosphate transport, the major species in the inner compartment, 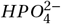, is transported from a compartment at pH 5 to a compartment at pH 7. At pH 5, the major species is however 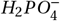, which loses one proton in the outer compartment when the transport occurs. As 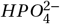 crosses the membrane with one proton, 20% of the species gain one additional proton from the inner compartment. The final equation is thus *Pi*_*e*_ *+* 0.2*H*_*c*_ *→ Pi*_*c*_ *+ H*_*e*_. c) Proton symport (transport of a metabolite coupled with a proton) of phosphate is the charge neutral transport of phosphate between an outer compartment at pH 5 and an inner compartment at pH 7.

Once the transported species and its protonation and charge state is identified, the final stoichiometry of the transport process can be determined. Because the protonation state of the metabolite may differ between compartments, protons can bind or release during transport (Fig. 3b). The equilibration of the transported species with the species distribution on each side of the membrane must be considered in the charge or proton balance (step 5). ‘Thermo-Flux’ calculates the protons that bind or release in each compartment based on the protonation state of the transported metabolite (step 2) and the pH in each compartment. In the previous example of phosphate transport, ‘Thermo-Flux’ compares the protonation state of phosphate in each compartment with the transported species and determines that, as the different species of phosphate equilibrate in the cytosol, one proton dissociates in the extracellular space and an additional 0.2 protons associate in the cytosol (Figure 3b).

If the specifically transported species has been experimentally determined, ‘Thermo-Flux’ offers the possibility to specify this particular species. ‘Thermo-Flux’ also allows the inclusion of several different transporter variants. Each variant represents the transport of a different species of the same metabolite, differing in protonation state and in net charge movement, and therefore potentially displaying a different thermodynamically feasible transport direction. Transporter variants are defined by coupling the transport of a metabolite to the transport of other metabolites such as protons or other ions, either in the same direction (symport) or in the opposite direction (antiport). For instance, variants can be added to include transport of a charge-neutral species of a metabolite at a given pH by co-translocation of protons together with the transported species to exactly balance the latter’s charge. Such transport variant ensures that for every metabolite, a transport variant exists that does not translocate net charge and therefore avoids over-constraining the model. For example, the major species of phosphate in the cytosol at pH 7, 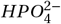, has two negative charges. Its transport from the extracellular space into the cytosol involves the movement of these negative charges. To add a charge-neutral transporter of phosphate from the extracellular space to the cytosol, a proton symport for phosphate transport (Figure 3c) can be added to the model. In the iMM904 case study, we added a charge-neutral variant for cytosolic and mitochondrial phosphate transporters using the code snippet 4.1 (Appendix A).

Some transport processes are more complex, and couple transport of a metabolite with chemical transformations such as in ATP powered proton pumps [34]. In general, these reactions are automatically processed and the transported species is correctly identified. However, in some cases the transported species is ambiguous, as either the same metabolite does not appear on both sides of the membrane, or protons pumped across the membrane cannot be distinguished from those being taken up or released by chemical transformations. In these ambiguous cases, ‘Thermo-Flux’ flags the reactions for manual curation. In the iMM904 case study, the transported compounds could be defined automatically for all reactions, except for (i) cross-membrane dehydrogenase reactions and for (ii) reaction mechanisms involving chemical transformations coupled to cross-membrane proton exchange. For instance, we had to specify the transported protons in the cytosolic ATPase (ATPS). In this reaction, a proton is pumped through the membrane during the chemical reaction, which we specified using code snippet 4.2 (Appendix A). For the reaction catalyzed by dolichyl phosphate mannosyl-transferase (DOLPMTcer), in which a chemical transformation occurs along with the transport, we had to manually specify the transported metabolite (code snippet 4.3, Appendix A).

### Step 5. pH-dependent charge and proton balancing

Most stoichiometric models do not contain accurate proton and charge balances. This is because the protonation and charge states of reactants are most often not consistently or accurately defined. Yet, charge and proton balances, as a function of specific compartmental pH values, are important to accurately describe the biochemical thermodynamics of a cell. To implement such balances, ‘Thermo-Flux’ first determines the net change in protons for each reaction, using the protonation state of every metabolite of this reaction as calculated previously (cf. step 2). To then balance the reaction, ‘Thermo-Flux’ compensates any excess proton by adding the equivalent amount of proton to the opposite side of the stoichiometric equation. For example, at pH 7 and ionic strength 250 mM, hydrolysing one ATP molecule releases 0.7 protons (Table 2). In this case, ‘Thermo-Flux’ balances the reaction by adding 0.7 protons to the right-hand side of the equation, which corrects charge imbalances that arise from differences in protonation states.

**Table 2.** Proton and charge balance of ATP hydrolysis at pH 7.

|  | ATP | + H<sub>2</sub>O | → ADP | + Pi | Net |
| --- | --- | --- | --- | --- | --- |
| Average protons | 12.1 | 2.0 | 12.2 | 1.2 | -0.7 |
| Average charge | -3.9 | 0.0 | -2.8 | -1.8 | -0.7 |

To capture the net charge movement of a transport process, we use a pseudo-metabolite, ‘charge’, representing the net positive charge of any metabolite crossing a membrane. When a transport process moves charge across the membrane, this metabolite is added to ensure charge balance (analogous to mass balance). To model the transport of a negative charge, we use the transport of a positive charge with opposite stoichiometry. As an example, in the uniport of phosphate from the outer compartment at pH 5 to the inner one at pH 7, the transported species is 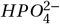 and carries two negative charges. Thus, this transport process is described by the reaction Pi_e_ + 0.2 H_e_ + 2 charge_c_ → Pi_c_ + 2 charge_e_ (Figure 3b), with the subscripts referring to the compartments (c for cytosol, e for extracellular). To balance the reactions of the iMM904 model in terms of protons and charges, we used the code snippet 5.1 (Appendix A).

Similarly to protons, Mg^2+^ ions can also be balanced. If magnesium ions are transported, they must be considered in the charge balance. However, note that magnesium affinity constants may not be available for all metabolites. Thus, if magnesium ions are balanced, we suggest allowing for free magnesium ion transport between compartments to prevent over-constraining the model to the magnesium ion balance. This can be done by adding a transport process for magnesium ions between each compartment, and not enforcing the second law of thermodynamics for these transport processes, such that they remain fully reversible.

### Step 6. Calculation of Gibbs energy of reactions

Next, we calculate the Gibbs energies of each reaction and transport process. With these Gibbs energies, we constrain the direction of the reactions and transport processes by means of the second law of thermodynamics (Figure 4a). In the following explanations, we refer to both transport processes and chemical transformations as ‘reactions’. In a cell, the Gibbs energy of a reaction (Δ_*r*_ *G*) differs from that under standard conditions. The Gibbs energy of reaction *j* is defined as,

**Figure 4.**
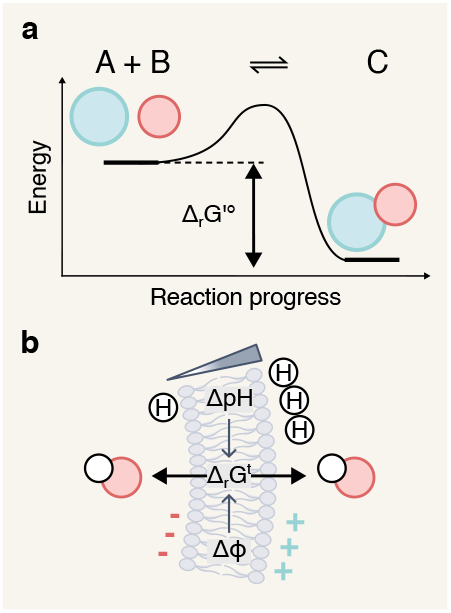
Calculation of Gibbs energy of reaction a) Under standard conditions, the Gibbs energy of reaction is the difference in energy between the product C and the reactants (A+B). (Δ_*r*_ *G*^*′°*^) is the standard Gibbs reaction energy transformed to the compartment’s physical and biochemical parameters (pH, ionic strength). b) The transport term of the Gibbs reaction energy for a transport process also depends on the cell’s parameters (differences in pH and in membrane potentials ΔΦ)

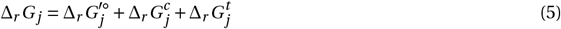

where 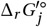 is the standard transformed Gibbs energy of the reaction, 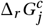 the Gibbs energy change due to the contribution of metabolite concentration, and 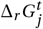 the Gibbs energy change due to metabolite transport.

The first term, 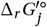, is calculated from the difference between the standard transformed Gibbs formation energy of products and reactants and an additional correction term to account for uncertainty in Gibbs reaction energies:

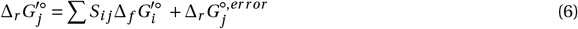

where *S*_*i j*_ is the stoichiometric coefficient of metabolite *i* in the reaction *j* and Δ_*r*_ *G*^*°,error*^ represents the uncertainty in Gibbs reaction energies. Because metabolites are shared among reactions, the uncertainty in Δ_*r*_ *G*^*°*^ must reflect the fact that the same metabolites participate in several reactions, meaning that these uncertainties are correlated. This correlation is captured using multivariate confidence intervals, computed with the square root covariance matrix of Gibbs reaction energies, *Q*, as returned by *eQuilibrator*, and an error vector *m*, which follows a normal distribution *m ∼ N* (0, 1), such that Δ_*r*_ *G*^*°,error*^ *=Q ·m*. This formulation ensures that the Δ_*r*_ *G*^*°,error*^ are internally consistent [22, 25, 19].

The second term of equation 5, Δ_*r*_ *G*^*c*^, accounts for the contribution of metabolite concentrations to the Gibbs energy of a reaction. This term is needed whenever metabolites are present at concentrations other than 1 M and can be calculated by,

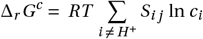

where *S*_*i j*_ is the stoichiometric coefficient of metabolite *i* in reaction *j* and *c*_*i*_ is the concentration of metabolite *i*. Note that concentrations can be used here instead of metabolite activities, because we use the extended Debye-Hückel theory in the calculation of Gibbs formation energies of metabolites (step 2).

Finally, the third term of equation 5, 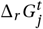, is specific to transport processes and represents the Gibbs energy change due to the movement of charge or protons across the membrane. It is influenced by (i) the transport of species between compartments with different pH values and the release or binding of protons caused by protonation or de-protonation of the transported species, by (ii) the translocation of additional free protons,e.g., by proton sym-/anti-porters or by proton pumps and (iii) the movement of charges between compartments with different chemical potentials. Taking these contributions into account, 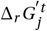, is defined by,

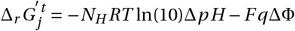

where *N*_*H*_ is the total number of protons transported across the membrane, Δ*p H* is the pH difference across the membrane, q is the number of charges transported across the membrane (determined in step 5), F is the Faraday constant and ΔΦ is the membrane potential difference.

### Step 7. Establishment of the thermodynamic-stoichiometric solution space

Initially, the flux solution space of the stoichiometric model we are converting includes the steady-state mass balance constraints (equation7) and the constraints on flux bounds (equation 8), typical of standard FBA :

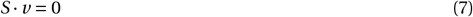

and each flux *v* _*j*_ of reaction *j* is within its lower and upper bounds *lb*_*j*_,*ub*_*j*_ :

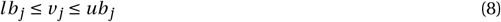

Having calculated the Gibbs reaction energy for all reactions, including transport processes, thermodynamic constraints can be added to all reactions (transport processes and chemical transformations) in the model. Importantly, the second law of thermodynamics constrains the direction of each reaction, such that only reactions with negative Gibbs reaction energy are allowed to proceed (equation 9), according to:

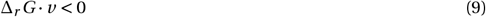

These ‘second law’ constraints prevent thermodynamically infeasible combinations of reaction fluxes, turning the solution space for the steady-state fluxes *v* into a non-convex space.

Furthermore, ‘Thermo-Flux’ also allows addition of a constraint on the cellular Gibbs energy dissipation rate, representing the sum of all reactions’ Gibbs reaction energy multiplied by the corresponding reaction flux, (see supplementary E). An upper limit in the cellular Gibbs energy dissipation rate has previously been shown to improve flux predictions, particularly those associated with overflow metabolism [21].

We used the code snippet 7.1 (Appendix A) to setup a FBA optimization problem with thermodynamic constraints of the iMM904 model. As in FBA, ‘Thermo-Flux’ allows for the selection of various objective functions for the optimizations, such as maximizing growth rate (biomass production), energy production or minimal ATP or redox potential consumption [13, 15]. Once the objective has been selected, the code snippet 7.2 (Appendix A) is used to optimize the model using the Gurobi solver [35].

#### Box 2

**Additional considerations for the formulation of the thermodynamic/stoichiometric constrained solution space**

**Metabolite concentration bounds**

Typically, intracellular metabolite concentrations vary between 0.1 *µ*M and 10 mM [16, 36], and this range is chosen as the default concentration range. For metabolites with concentration measurements reported in the literature, these bounds can be adjusted to describe the minimal and maximal reported values.

**Some metabolite concentrations can be expressed as ratios**

For metabolites that always appear in pairs and for which there are no explicit biosynthesis reactions in the model (e.g., redox cofactors such as NAD(H) and NADP(H)), using the ratio of their concentrations may be preferred over using absolute concentrations. In such cases, the concentration of one of the forms is fixed to 1 M and the other represents the ratio, which can also be constrained with an upper and a lower bound.

**Compartment-specific metabolite concentrations**

The experimental determination of metabolite concentrations in distinct subcellular compartments is experimentally challenging [37, 38]. In contrast, whole-cell concentrations are readily available from metabolomics studies. Importantly, whole-cell concentration data can be expressed in terms of compartmental concentrations if the relative compartmental volumes are known. The whole-cell concentration of a metabolite is a weighted average of the concentration in the different subcellular compartments:

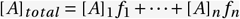

where [*A*]_*total*_ is the concentration of a metabolite in the whole cell, [*A*]_*n*_ is the concentration of the metabolite in compartment *n* and *f*_*n*_ is the fractional volume of compartment *n*. Whole-cell concentration measurements can be used as additional bounds during the FBA optimization (step 7) or in regressions.

**Metabolites for which concentration are ignored**

The concentration of the pseudo-metabolite ‘charge’ and of other pseudo-metabolites (e.g. biomass) are ignored when calculating the Gibbs energy term accounting for the contribution of metabolite concentrations, Δ_*r*_ *G*^*c*^. The concentration of protons is also ignored in this term, as protons are already considered in the pH-corrected transformed Gibbs energies of formation (step 3).

**Reactions where the second law of thermodynamics is not applied**

The second law constraint is not applied for exchange reactions, where metabolites are exchanged across the system boundary. This constraint also is not applied to the transport of water between compartments, which is assumed to be fully reversible. To ignore the second law constraint for the transport of water, the code snippet B2 (Appendix A) can be used.

#### Box 3

**Regression — fitting models to experimental data**

Through a regression analysis, models can be fit to available data (e.g. metabolite concentrations and/or extracellular fluxes), enforcing simultaneous compliance with the laws of thermodynamics and experimental measurements and enabling the accurate inference of non-measured parameters (e.g., Δ_*r*_ *G*^*°,error*^). Furthermore, the cellular Gibbs energy dissipation rate can also be inferred from regression to experimental data [21]. In ‘Thermo-Flux’, the code snippet B3 (Appendix A) can be used to add regression-specific variables (e.g., whole-cell concentrations) and objectives to the previously constructed linear program with thermodynamic and stoichiometric constraints.

A linear optimization is used to find the minimum of an objective function that quantifies the difference between predicted and experimental values of metabolite concentrations, fluxes or both. These regressions can be performed simultaneously on datasets of multiple experimental conditions, such as those obtained from cells grown on different substrates. These simultaneous regressions could help determine more accurate values of Δ_*r*_ *G*^*°,error*^, which would be consistent between datasets. The need for such improved Δ_*r*_ *G*^*°*^ estimates is strong for reactions in which the component contribution method only provides highly uncertain Δ_*r*_ *G*^*°*^ (step 3).

## 4 Results

Having developed the pipeline to automate the conversion of stoichiometric models into combined thermodynamic-stoichiometric models, our first goal was to assess the pipeline’s capacity to do so. To this end, we obtained all 107 stoichiometric models, reflecting various organisms, from the BiGG database [39] and ran them through our pipeline. ‘Thermo-Flux’ automatically performed the full conversion, up to the establishment of the thermodynamic-stoichiometric solution space (all steps), for 87 models (77% of all available models). Among the 20 models that could not be automatically processed, we identified three main shortcomings (Figure 6a). First, seven models had initialization errors. Specifically, the loading of four models failed due to missing biomass equations or incomplete model annotations (e.g., missing compartment localization), and three yielded infeasible FBA simulations, based on the original stoichiometry alone. Then, six other models had transport or biomass reactions that could not be handled because they involved more than three compartments. Currently, ‘Thermo-Flux’ only handles reactions involving up to three compartments, as dealing with a larger number poses challenges in identifying the transported metabolites (see supplementary H). Finally, in seven additional models, automatic balancing failed because of reactions that involved a chemical transformation during a transport process. For example, transport reactions via the phosphotransferase system could not be balanced automatically in some *E. coli* models. Automatic balancing also fails when models contain metabolites with undefined or ambiguous structure or formula, which happened mostly for specific redox cofactors, such as ferricytochrome or electron-transferring flavoproteins. Overall, the high success rate shows that ‘Thermo-Flux’ can facilitate the automated conversion of stoichiometric models into combined thermodynamic-stoichiometric ones.

**Figure 5.**
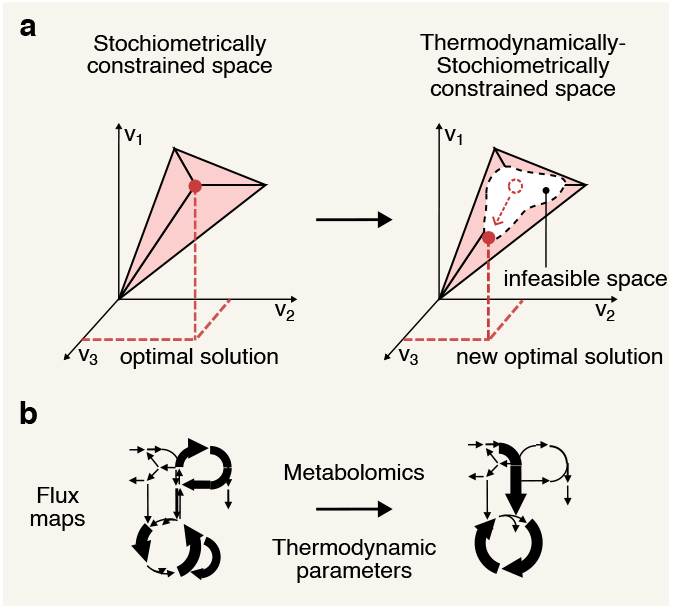
Thermodynamic-stoichiometric solution space. a) The addition of the second law to the stoichiometrically constrained solution space removes thermodynamically infeasible combinations of reactions, turning the flux cone into a non-convex space and potentially altering the optimal solution. b) By adding thermo-dynamic constraints and incorporating experimental data, such as metabolite concentrations, more accurate flux predictions can be obtained.

**Figure 6.**
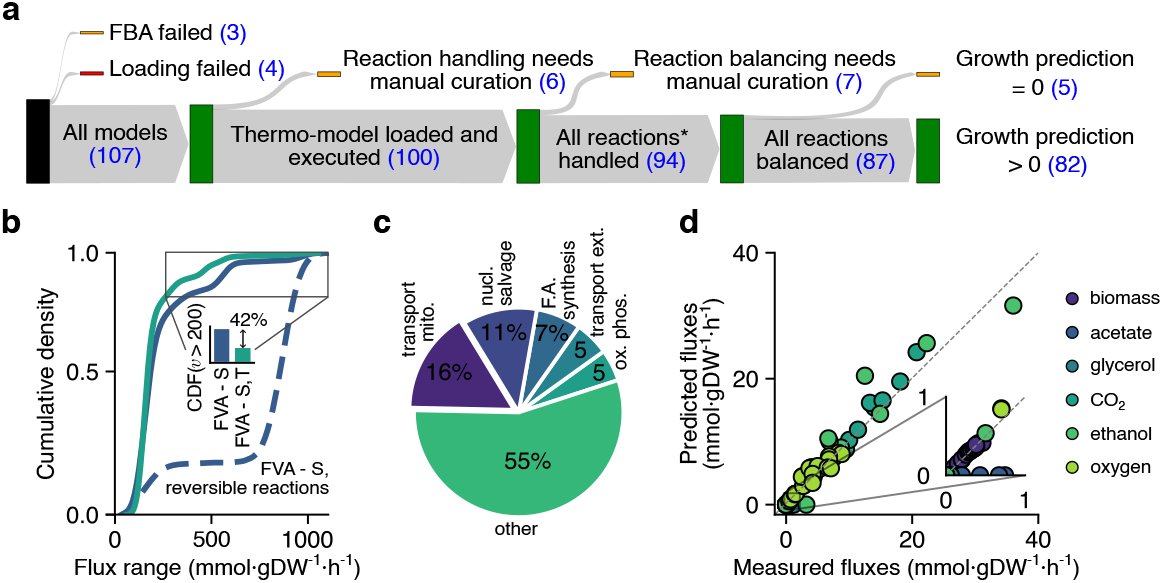
a) Application of the ‘Thermo-Flux’ pipeline to all 107 models from the BiGG database, with the number of models that passed each automated step of the pipeline. b) Cumulative density distribution of flux ranges assessed using Flux Variability Analysis with thermodynamic constraints (FVA - S,T), Flux variability Analysis while keeping the predefined reaction directions (FVA - S) and FVA when considering all reactions as reversible (FVA - S, reversible reactions). Reactions with a flux range above 200 mmol·gDW^*−*1^·h^*−*1^ decreased by 42% when performing (FVA - S,T) relative to (FVA - S), both with predefined reaction directions, as indicated by the inset bar plot. c) Pathways for which thermodynamic constraints alter reaction directionality relative to standard FBA (FBA - S), in at least one of the 20 glucose uptake rates for which simulations were performed (from 0 to 20 mmol·gDW^*−*1^·h^*−*1^). Percentages indicate the proportion of such reactions within each pathway. Pathways contributing less than 5% are grouped into the ‘other’ category. d) Predictions of extracellular fluxes using (FBA - S,T) are compared with experimentally measured fluxes for different conditions. For each condition, we set bounds on glucose uptake rates for the different conditions and maximized growth rate as an objective. Additionally, we set a constraint on the upper limit of the Gibbs dissipation rate. The experimentally measured fluxes were taken from [40, 41, 42].

We then established how many of the 87 converted models would allow for biomass synthesis while having reaction directions constrained by the second law of thermodynamics. For this, we added thermodynamic constraints (step 7), default physical and biochemical parameters (step 1), and broad ranges of metabolite concentrations (box 3), and defined the objective function as maximization of growth rate. We found that 82 models displayed growth (i.e. a growth rate above zero), showing that the addition of thermodynamic constraints did not prevent essential reactions from occurring in the direction necessary for biomass formation. On the other hand, the remaining five models, which had displayed growth rates above zero prior to the addition of these constraints, no longer displayed any growth. This observation indicated that essential reactions were blocked by the imposed thermodynamic constraints. One possibility for this to occur could be that the default physical and biochemical parameters and metabolite concentration ranges are incompatible with the essential metabolic reactions having a certain required flux in these models. Another possibility is that constraining the reaction directions would require additional reactions, such as transporter variants, to be defined. Overall, most models generated by ‘Thermo-Flux’ could be used to perform flux predictions.

Next, using the iMM904 yeast model as a case study, we asked to what extent augmenting a stoichiometric model by adding comprehensive proton and charge balances and thermodynamic constraints with ‘Thermo-Flux’ would narrow down the flux space and improve the flux predictions. To test this, we examined three distinct aspects. First, using flux variability analysis, we compared the flux ranges of the iMM904 stoichiometric model using fully reversible reactions (FVA - S, reversible reactions) with the flux ranges of the same model, but keeping reaction directions as defined in the BiGG model (FVA - S). Here, we found that predefined reaction directions significantly constrained the flux ranges, as seen from the difference between the flux ranges of the respective model variants (Figure 6b). Then, when determining the flux ranges using flux variability analysis on a thermodynamically-constrained model (FVA - S,T) with predefined reaction directions, we found that the number of reactions with unrealistically large flux ranges decreased by 42% compared to the stoichiometric model with predefined reaction directions (FVA - S) (Figure 6b).

Secondly, we investigated the extent to which the steady-state flux directions are modified in the thermodynamic-stoichiometric model. To do so, we performed FBA with thermodynamic constraints (FBA - S,T) simulations over a range of glucose uptake rates (GURs) and compared the resulting reaction directions with those obtained from standard FBA (FBA - S) simulations, both with the same predefined directions. We found that, across all GURs, the direction of 15% of all reactions had been altered in thermodynamically-constrained (FBA - S,T) solutions, especially in mitochondrial transport processes and metabolic reactions involved in nucleotide salvage and fatty acid synthesis (Figure 6c). Thus, important reactions for which the model did not provide predefined directions were constrained further in the thermodynamically-constrained (FBA - S,T) solutions.

Finally, we asked whether flux predictions from thermodynamically-constrained optimizations (FBA - S,T) would quantitatively agree with experimental data. Importantly, the (FBA - S,T) simulations were additionally constrained using an upper limit on the cellular Gibbs dissipation rate [21]. Here, we found that predictions of extracellular fluxes over multiple conditions spanning a wide range of GURs agreed with experimental data, including *CO*_2_ and *O*_2_ exchange fluxes and ethanol secretion (Fig 6d).

Thus, we showed that augmenting a stoichiometric model with charge and proton balancing, considering pH and charge gradients in cross-membrane transport and adding thermodynamic laws impacts (i) flux ranges by narrowing them and removing thermodynamically-infeasible loops and (ii) directions of important reactions linked with energy generation and central metabolism (Fig 6c). We also showed that (iii) the augmented model could accurately predict yeast physiology (Fig 6d).

## 5 Discussion

To facilitate the use of thermodynamic constraints in metabolic modeling, we developed the ‘Thermo-Flux’ pipeline to convert stoichiometric metabolic network models into combined thermodynamic-stoichiometric models, in fully- or semi-automated manner. The workflow of this pipeline includes the incorporation of physical and biochemical parameters of cellular compartments, automated definition of transporter variants and calculation of membrane transport Gibbs reaction energy, accurate addition of proton and charge balances, and addition of thermodynamic constraints for all reactions and transport processes. ‘Thermo-Flux’ could automatically process 77% of the models in the BiGG database (ranging from bacteria to eukaryotic cells) and convert them into combined thermodynamic-stoichiometric models. While it failed for a minority of models for reasons mentioned above, we have demonstrated with the iMM904 model that with limited additional manual effort, ‘Thermo-Flux’ can convert large stoichiometric models into combined thermodynamic-stoichiometric models that enable improved predictions of metabolic fluxes.

There are several aspects to consider when predicting metabolic fluxes with thermodynamic-stoichiometric models. First, these models are sensitive to physical and biochemical parameters such as pH and membrane potential differences. These values dictate (i) the direction of proton and charge transport across membranes. As transport processes are an essential part of stoichiometric networks, correct directionality for these processes is crucial to obtain valid flux predictions. Furthermore, stoichiometric networks often lack predefined directions for many transport processes [43], therefore these processes can only be constrained further using thermodynamic-stoichiometric models (Fig. 6c). Physical and biochemical parameters also determine (ii) the amount of protons (and charge) involved in each reaction, influencing the role of various pathways in proton- and charge balancing [44, 45]. Therefore, it is important to use organism-specific values for these parameters.

Second, thermodynamic-stoichiometric models also require definitions for upper and lower bounds on metabolite concentrations. As such bounds together with the second law of thermodynamics define reaction directionality, *a priori* assumptions on reaction directions are not needed anymore in thermodynamic-stoichiometric models. Instead, users can define broad metabolite concentration ranges, e.g. from 0.1 µM to 10 mM, or narrower ranges based on metabolome measurements. The narrower the metabolite concentration ranges, the more constrained the flux space will be, which will translate into better flux predictions— provided that the measurements used are accurate, otherwise predictions will be incorrect as well. Advances in metabolomics now allow quantitative measurement of whole-cell concentrations of an increasing number of metabolites [46, 40, 47], and these measurements could be valuable to constrain reaction directions in a condition-specific manner.

Looking ahead, we anticipate that models built with ‘Thermo-Flux’ will advance both fundamental and applied research. On the fundamental side, models augmented with ‘Thermo-Flux’ can address several challenges: First, Gibbs energies estimated by the component-contribution method still contain uncertainties. Through fitting thermodynamic-stoichiometric models to extracellular fluxes and metabolome data, it should be possible to further narrow down such uncertainties and to adjust standard Gibbs energies to *in vivo* conditions. Second, by explicitly modeling dependence to physical and biochemical parameters (pH, ionic strength, temperature, and membrane potential) models augmented with ‘Thermo-Flux’ rigorously link flux predictions to cellular conditions and thus allow systematic sensitivity analyses to physical and biochemical parameters. Enabling, for example, investigation of energetic/mechanistic features of proton- and charge-coupled pathways, as done in [44, 45]. Third, models developed with ‘Thermo-Flux’ facilitate the analysis of thermodynamic principles on the genome-scale, from limits on growth and candidate cellular objectives, to core relationships such as the trade-off between yield and growth rate [48, 49], upper bounds on Gibbs-energy dissipation [21], the flux–force relationship [50], and the max–min driving force (MDF)[51, 18]. On the applied side, thermodynamic-stoichiometric models developed with Thermo-Flux’ can support biotechnology and metabolic engineering through improved predictive capabilities [52, 20] and the possibility of uncovering thermodynamic bottlenecks [19].

## 6 Methods

### Applying ‘Thermo-Flux’ on all 107 models from the BiGG database

We downloaded every model in the BiGG database via the BiGG Models web API. According to the database version information, the API version was “v2” and was last updated on the 31/10/2019. The BiGG models version was “1.6.0”. We downloaded the models in the “.xml” format. We used ‘Thermo-Flux’ to load every model and convert it to a thermodynamic-stoichiometric model. We then set the objective to the biomass exchange reaction, erased all prior reaction directions, and first optimized the models using Cobra (for a stoichiometric optimization only).

While applying the ‘Thermo-Flux’ pipeline, we designed several checks to ensure that each model could go on to the next step. The first verification was making sure that the model could be read and executed by the cobra loading and optimizing functions. Then, we selected models with only reactions that can be handled by ‘Thermo-Flux’ (biomass reactions that were not handled were not counted). We further restricted our list of models by keeping only the ones that had all their reactions successively balanced by ‘Thermo-Flux’. Finally, the thermodynamically-constrained FBA(FBA S,T) growth rate predictions were obtained using the Gurobi model set up with `tmodel.add_TFBA_variables()`. Growth rate predictions lower than 1e-6 were not considered as positive growth prediction but rather coming from numerical errors. The calculations for (FBA - S,T) were run on the university’s computational cluster “Hábrók” and were optimized until the optimal solution was found, with a maximal duration of 24 h each.

### Calculating the flux ranges for the iMM904 model

First, we performed a variability analysis to compute the flux bounds for the iMM904 model with its predefined reaction directions set as in the BiGG database. Then, we set all flux bounds to (-500,500) and performed a variability analysis again. Both calculations were done using the cobrapy function:

`flux_analysis.flux_variability_analysis()`

After having augmented iMM904 into a comprehensive stoichiometric-thermodynamic model with ‘ThermoFlux’ (the conversion steps are shown in Appendix A), we performed variability analysis as with the function implemented in ‘Thermo-Flux’:

`solver.gurobi.variability_analysis()`

We used this function to set an optimization problem for fluxes. Specifically, the function uses the Gurobi multi-scenario optimization feature, with two scenarios for each flux (one minimizes the flux and the other maximizes it). The results were retrieved with:

`solver.gurobi.variability_results()`

Finally, to identify reactions with an inflated flux range, we calculated the number of reactions with a flux range above the threshold of 200 mmol·gDW^*−*1^·h^*−*1^ for both the augmented iMM904 model and the iMM904 model with predefined reaction directions.

### Simulations at different glucose uptake rates

To study the impact of thermodynamic augmentation on steady-state flux directions, FBA with thermodynamic constraints (FBA - S,T) and FBA using only stoichiometric constraints (FBA - S) simulations were conducted on the iMM904 model across a range of glucose uptake rates (GURs). For each GUR (0.5 to 20 mmol·gDW^*−*1^·h^*−*1^), we performed both (FBA - S) and (FBA - S,T) on the augmented model. All simulations were performed with growth rate (biomass_EX flux) maximization as the objective, using the ‘Thermo-Flux’ function : `solver.gurobi.variable_scan()`

The direction of each reaction was classified as forward, backward, or inactive, based on its flux sign in the optimal solution. Reaction directionality changes between the (FBA - S) and (FBA - S,T) solutions was calculated as the fraction of reactions that changed direction in any of the simulated GUR conditions over the total number of reactions. We classified reactions according to their pathways using the Subsystem annotations from the BiGG database.

### Comparison of thermodynamically-constrained predictions and extracellular reaction fluxes

To evaluate the flux predictions from (FBA - S,T) simulations over a range of different physiological condition for *S. cerevisiae*, we ran multiple (FBA - S,T) optimizations. Specifically, for each different condition, we set the glucose uptake rate of the model to the measured one and compared the remaining predicted extracellular fluxes with the measured reaction rates. Furthermore, the cellular Gibbs energy dissipation rate was constrained by 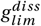 as taken from [21]. We set the maximization of growth rate as an objective. The experimental data used for comparison with the model predictions were compiled from several published datasets, including (i) *S. cerevisiae* CEN.PK strain grown in glucose-limited chemostat cultures at different dilution rates [40], DS2891X strain grown in glucose-limited chemostat cultures at different dilution rates and in one batch culture[41], and (iii) data from a batch-cultured glucose grown CEN.PK strain [42].

## Acknowledgments

Funding is acknowledged from the Dutch Research Council (NWO) (VICI, VI.C.192.003, to MH), and from the European Union’s Horizon 2020 research and innovation programme (Gain4Crops, grant agreement 862087, to MH). We thank the Center for Information Technology of the University of Groningen for their support and for providing access to the Hábrók high performance computing cluster; Yusuke Himeoka, Moritz Beber, Elad Noor and Mattia Gollub for helpful discussions and suggestions.

## Appendices

## A Appendix Code Snippets

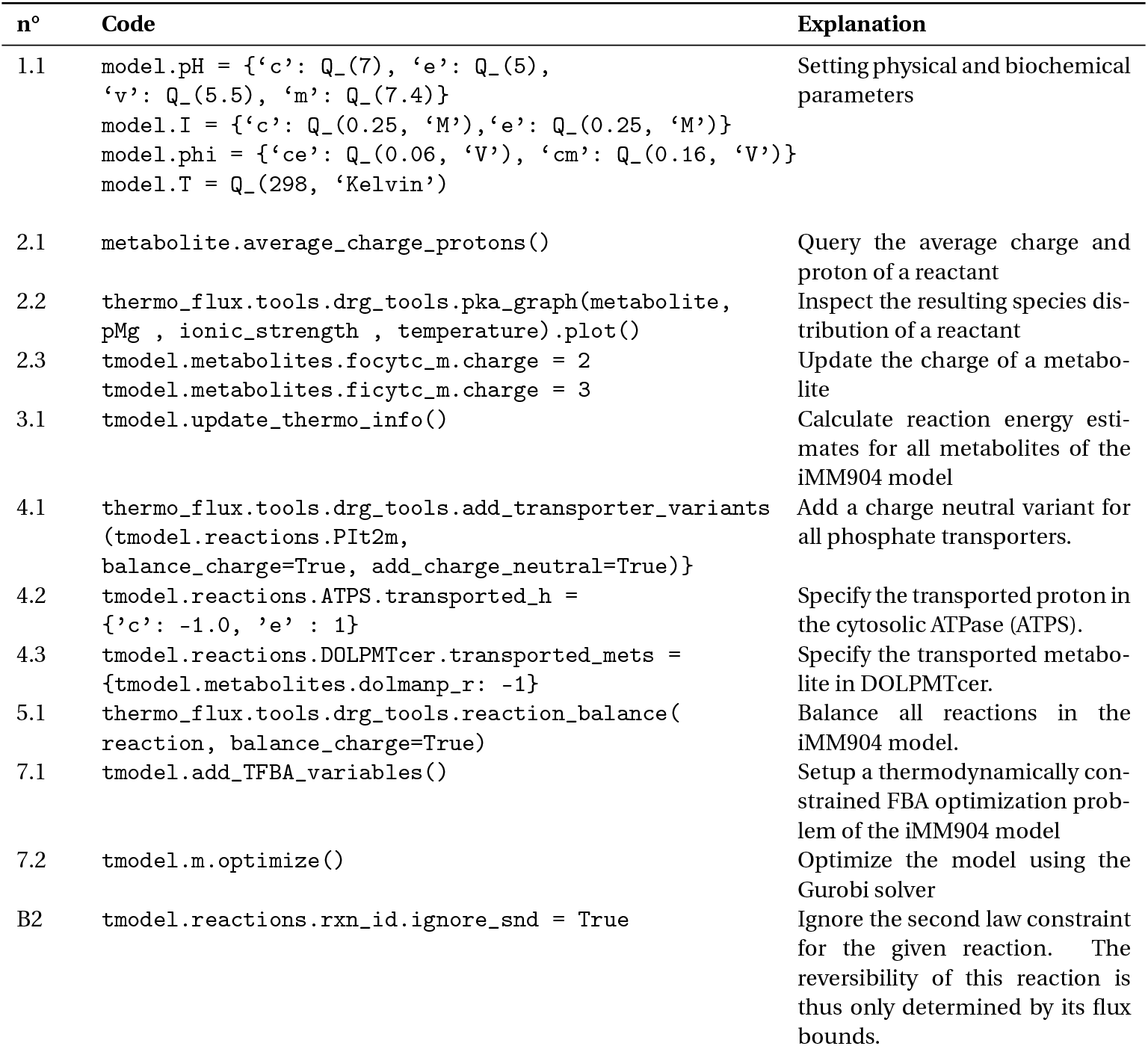
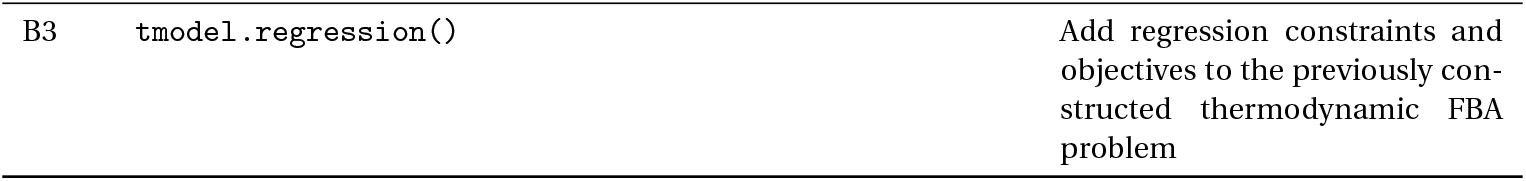

## Supplementary information

### B Example physical and biochemical parameters

**Table S1.** Example physical and biochemical parameters of *E. coli, S. cerevisiae*, and *A. thaliana* leaf compartments.

| pH |  |  |  |
| --- | --- | --- | --- |
| Compartment | *E. coli* | *S. cerevisiae* | *A. thaliana* leaf |
| Cytosol | 7.6 (Shimamoto et al., 1994) | 7.0 (Orij et al., 2009) | 7.3 |
| Extracellular/apoplast | 7 (Shimamoto et al., 1994) | 5.0 (Weller et al., 2005) | 4.7 |
| Mitochondria | – | 7.4 (Orij et al., 2009) | 8.1 |
| Vacuole | – | – | 6 |
| Plastid stroma | – | – | 7.3 – 7.6 |
| Plastid Thylakoid | – | – | 7.5 – 5.7 |
| Peroxisome | – | – | 7.5 |
| Golgi | – | – | 6.9 |

| Membrane potential (mV) |  |  |  |  |
| --- | --- | --- | --- | --- |
| Compartment inside | Compartment outside | *E. coli* | *S. cerevisiae* | *A. thaliana* leaf |
| Cytosol | Extracellular | -150 | -60 | -150 |
| Mitochondria | Cytosol | – | -160 | +135 – +170 |
| Vacuole | Cytosol | – | – | -80 – -20 |
| Plastid stroma | Cytosol | – | – | +111 – +123 |

| Compartmental relative volume (%) |  |  |
| --- | --- | --- |
| Compartment | *S. cerevisiae* | *A. thaliana* leaf |
| Cytosol | 90–95 (Canelas et al., 2011) | 3.8 (Tollet et al., 2024) |
| Mitochondria | 5–10 (Canelas et al., 2011) | 2.1 (Tollet et al., 2024) |
| Vacuole | – | 84.3 (Tollet et al., 2024) |
| Plastid | – | 9.4 (Tollet et al., 2024) |
| Peroxisome | – | 0.09 (Tollet et al., 2024) |
| Golgi | – | 0.009 (Tollet et al., 2024) |
| ER | – | 0.1 (Tollet et al., 2024) |

### C Description of the typical model and constraints

**Table S2.**
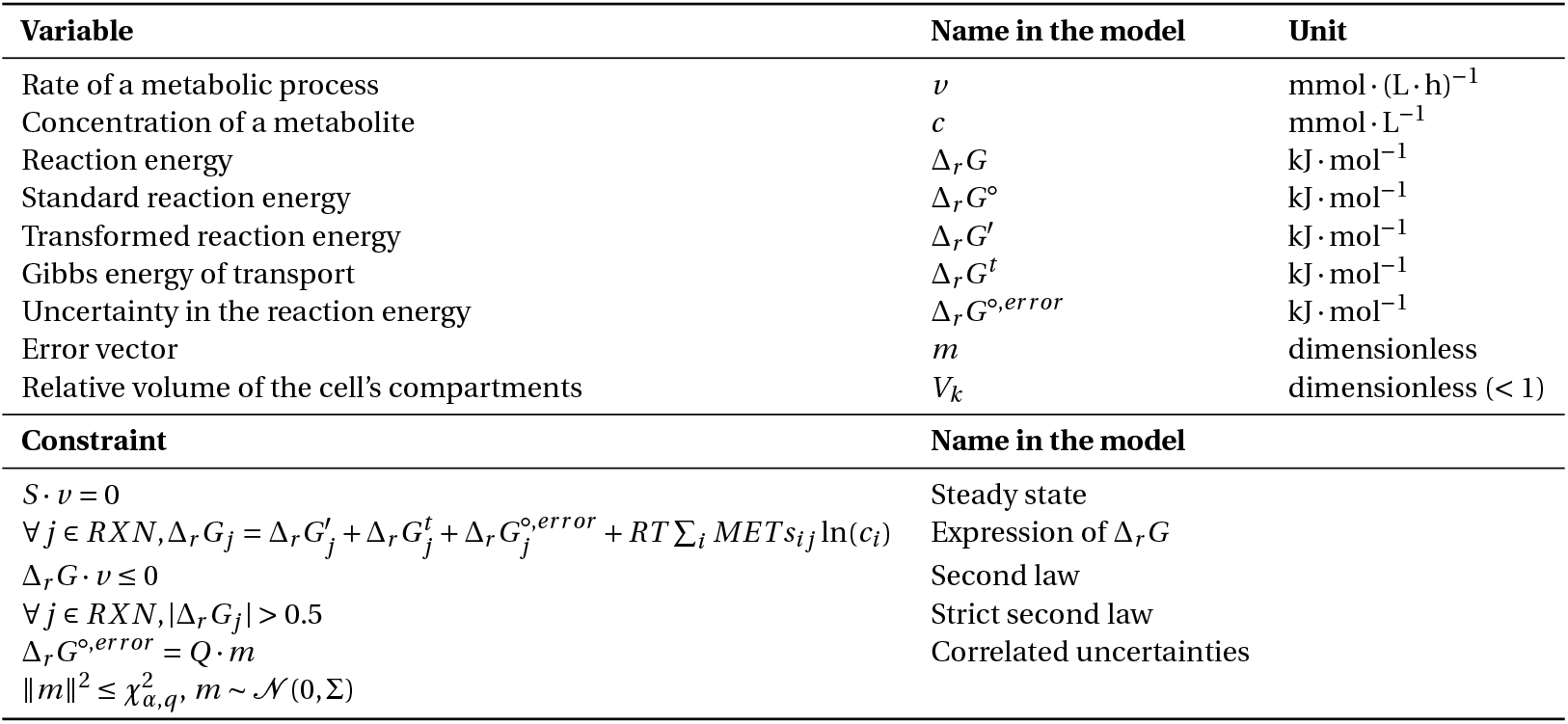
Variables and constraints in the metabolic model. *χ*_*α,q*_ is the *α*-quantile of the *χ*^2^ distribution with *q* degrees of freedom.

### D Implementing conditional constraints in a linear program

The second law constraint relies on a conditional statement over the sign of the flux *v* that we consider. In linear programming, such constraints are implemented by introducing an auxiliary binary variable *b* as the indicator of the sign.

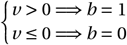

Therefore, the sign of the flux *v* depends on the value of the binary variable, which is enforced by implementing the following constraints.

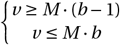

The big-M value should be chosen as tight as possible to avoid numerical issues [35]. Setting equal to the largest of the upper bounds of *v* often works well.

To eventually model the second law, we apply the same logic on the Δ_*r*_ *G* variable.

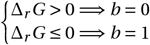

As done previously, this conditional constraint is enforced by:

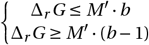

where M’ is this time bigger than the maximum possible value of Δ_*r*_ *G*.

### E Gibbs energy dissipation rate

The Gibss energy dissipation rate is the sum over all exchange reaction of the product of the reaction energy and the rate at which this reaction is going. It is implemented by defining a new variable `g_2` that equals the latter sum and setting an upper bound to `g_2` [21].

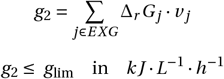

### F Biomass formation energy

Firstly, the standard enthalpy of combustion 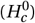 can be calculated using the Patel-Erickson equation (eqn. 1) which assumes it is proportional to the number of electrons that are transferred to oxygen during combustion where n_C_, n_H_, n_°_, n_N_, n_P_ and n_S_ are the number of C, H, O, N, P and S atoms in the biomass empirical formula [53, 30].

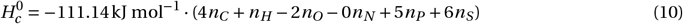

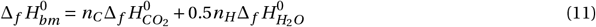

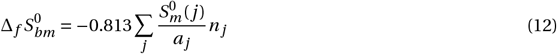

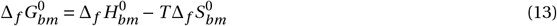

Molar biomass Gibbs formation energy can be converted to specific formation energy using the density of cell dry weight or the molecular mass calculated form the empirical formula.

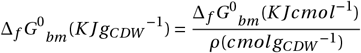

### G Units of the usual fluxes and reaction energy compared to the units of biomass-related values

When specifying the value of the formation energy of the biomass metabolite, we should plug in the values in units of J/gDW. This differs from all the other reactions in the model, for which values of kJ/mol are used.

The reason for this discrepancy has to do with the units of the fluxes. Indeed, the biomass reaction is often based on cellular composition measurements, which give the cellular content in grams per dried gram of cell (g/gdW).

For a normal reaction:

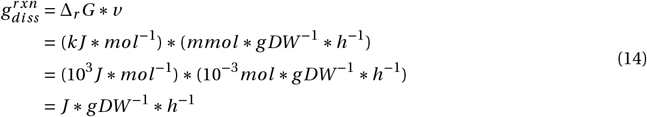

For the biomass reaction:

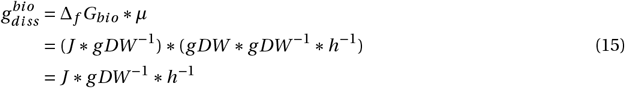

Since thermoflux handles all reactions the same when determining the gibbs energy dissipation rate (multiplying *v* by add Δ_*r*_ *G*), we have to pretend that the biomass formation energy has units of (*k J *mol*^*−*1^).

### H Balancing three-compartment transport reactions

The tool developed in the package works on transport reactions between exactly two compartments. The case of reactions that involve three compartments, for example reactions that use the TON motor complex (in Gram-negative bacteria) [54], require a few additional steps than in the regular case. When a three-compartment reaction is given to the balancing function, the latter identifies the sub-transport reactions that compose the total reaction. This is achieved by spotting the metabolites that travel from one compartment to another without being transformed (practically, if the metabolite’s identifier without the compartment annotation is untouched). After the reaction has been decomposed between sub-transport reactions and one core reaction with removed metabolites (i.e., metabolites present in the sub-transport reactions are taken out), the code can balance each entity separately. This approach only works if the core reaction is left with a maximum of two compartments, therefore this approach does not work for (i) reactions with transported metabolites that are converted during transport, because the core reaction could be left with more than 2 compartments and (ii) for reactions that involve more than three compartments. This approach has not yet been extended to transport reactions with more than three compartments.

### I Metabolites with unknown formation energy

By default, the mean formation energy of a metabolite with an unknown or non-decomposable structure is 0*k J*.*mol*^*−*1^. Therefore, formation energies of such compounds are not considered when calculating the reaction energies. This can cause issues in downstream calculation of reaction energies by requiring large absolute errors to estimate reasonable or feasible reaction energies. To obtain more reasonable estimates for completely unknown formation energies the information within the model reaction stoichiometry can be used.

Metabolites with completely unknown formation energy are present in reactions that contain other metabolites with known formation energy. For reactions where an unknown compound is present, the reaction energy is shifted by as much as the unknown formation energy. The better the estimate of the unknow formation energy, the closer the reaction energy is from its real value. Indeed, as systems tend to operate near equilibrium, reactions energies should be much closer to 0 than formation energies. We can thus assume that the current Gibbs standard energy is largely due to the gap between the real formation energy of the unknown compound and its current computed value – which is 0. Bringing the computed formation energy towards cancelling the current reaction energy will reduce the latter difference and lead to a better estimation of the formation energy of unknow compounds. The correlation between reactions is conserved as the minimization problem can be solved while taking multiple equations into consideration.

In practice this problem is implemented by regressing – for reactions that include unknown compounds - the uncertainty of reaction energies to the opposite of the set of mean standard reaction energies. The least-squares solution is an error vector, m, that can be multiplied by the square root covariance matrix (the part that relates to unknown compounds). This latter conversion in formation energy enables the definition of the new mean formation energies of the unknown compounds.

Using ATP hydrolysis as an example say the formation energy of ATP was unknown and we assume it to be 0 *k J*.*mol*^*−*1^. The Δ_*r*_ *G* of ATP hydrolysis then becomes -2802 *k J*.*mol*^*−*1^.

| Metabolite | Stoichiometry | Formation energy × Stoichiometry |
| --- | --- | --- |
| ATP | -1 | 0 |
| H<sub>2</sub>O | -1 | 238 |
| ADP | +1 | -1945 |
| Pi | +1 | -1095 |
| **Total (reaction energy)** |  | **-2802** |

If we now try to shift the formation energy of ATP to be as close as possible to the reaction energy of this reaction then it will approach a value of -2802 *k J*.*mol*^*−*1^ which is relatively close to the true formation energy of -2811 *k J*.*mol*^*−*1^ or at least a better estimate than 0 *k J*.*mol*^*−*1^. The more reactions a metabolite is involved in and the more known metabolites are in those reactions the better the estimate should be. In contrast, if multiple metabolites in a reaction are unknown then the estimated formation energy will be relatively further from the true value.

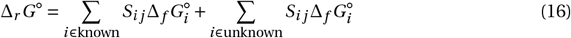

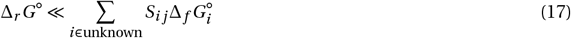

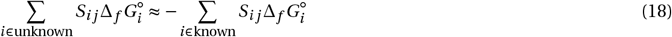

## Notes

### Competing Interest Statement

The authors have declared no competing interest.

